# Innate sleep apnoea in spontaneously hypertensive rats is associated with microvascular rarefaction and neuronal loss in the preBötzinger Complex

**DOI:** 10.1101/2023.08.09.552638

**Authors:** Reno Roberts, Robert T. R. Huckstepp

## Abstract

Sleep apnoea is a major threat to physical health and carries a significant economic burden. These impacts are worsened by its interaction with, and induction of, its comorbidities. Sleep apnoea holds a bidirectional relationship with hypertension which drives atherosclerosis, ultimately culminating in vascular dementia. To enable a better understanding of this sequalae of events, we investigated innate sleep apnoea in spontaneously hypertensive rats (SHRs), which have a range of cardiovascular disorders. SHRs displayed a higher degree of sleep disordered breathing, which emanates from poor vascular health leading to a loss of preBötC neurons. This model also displays small vessel white matter disease, a form of vascular dementia, likely associated with neuroinflammation in the hippocampus and the related deficits in both long- and short-term memory. Therefore, hypertension induces sleep apnoea through vascular damage in the respiratory column, culminating in neuronal loss in the inspiratory oscillator. This induction of sleep apnoea which in turn will independently exacerbate hypertension and neural inflammation, increasing the rate of vascular dementia.

## Introduction

Sleep apnoea (SA, >10s of respiratory depression) is estimated to effect 1.5 million UK adults, equivalent to ∼4% of men and ∼2% of women in the UK (Young et al., 1993). However, up to 85% more cases are undiagnosed [1], meaning the actual prevalence is closer ∼50% of men and ∼25% of women as seen in other large cohort studies in northern Europe [2]. This rise may be in part due to factors such as obesity and hypertension [3, 4], which are steadily rising in many developing countries [5].

Dyspnoeic episodes result in temporary but immediate elevations in blood pressure. The associated blood oxygen desaturation coupled with prolonged sympathetic activation may cause sustained hypertension [6]. As greater rates of dyspnoeic episodes lead to hypertension [6], and increasing severity of SA associated with linear increases in blood pressure [7], sleep apnoea has long been considered a risk factor for hypertension. Though it must be noted the association between hypertension and sleep apnoea may be age dependent [8].

Hypertension is a major risk factor to the world’s most prevalent killers, with the most emergent including coronary heart disease (CHD) and stroke [9]. Moreover, hypertension [10, 11] and sleep apnoea [12-14] are leading causes of Vascular Dementia (VaD), the second most common type of dementia worldwide [15]. The resulting release of pro-inflammatory cytokines, such as interleukin-6, caused by periods of intermittent hypoxic hypercapnia during sleep apnoea undoubtedly increases structural damage to endothelial cells of blood vessels in the brain [16], increasing the risk of small vessel white matter disease, a contributing factor to VaD [17, 18]. Vascular cognitive impairment (VCI), a complication of cerebrovascular disease [19], involves a wide range of disorders, the most severe of which is VaD. These cerebrovascular pathologies are more widespread in neurodegenerative disease with cognitive decline, as 20%-40% of dementia diagnoses are associated with VCI, most of which are linked directly to hypertension as a risk factor for developing the disease in mid-life [20].

Spontaneously hypertensive rats (SHRs) display cognitive deficit due to small vessel white matter disease, a form of vascular dementia [21]. SHRs develop cardiovascular pathologies that correlate with the presentation and progression of hypertension as is seen clinically [22]. SHRs also display sleep apnoea very early on in their disease progression, which can be blocked if blood pressure is normalised pharmacologically by systemic administration of the vasodilator hydralazine, a commonly used blood pressure medication [23]. This provides very clear evidence that hypertension may be an independent risk factor for sleep apnoea. Whilst hypertension reduces airway patency in a way that could induce sleep apnoea [24-26], we believe other mechanisms may contribute. We postulate that microvascular rarefaction also occurs in the preBötC of SHRs, leading to cell death and ultimately sleep apnoea. The sleep apnoea will then drive, or exacerbate, hypertension, microvascular rarefaction, and neural inflammation to cause cognitive decline and eventually dementia. Given the relationship between sleep apnoea and hypertension this will drive a downward spiral amplifying vascular dementia in these rats.

## Methods

### Surgeries

Adult (6-8 months old) male Wistar Kyoto rats (WKY: 417 ± 35g, *n =* 9) and Spontaneously Hypertensive rats (SHR: 406 ± 54g, *n =* 9)). The order of the rats pseudorandomized prior to surgery. Five animals were excluded as they did not complete the study.

Rats were induced and maintained on 0.5-2% inhalation Isoflurane (Piramal Healthcare, Mumbai, India) in pure oxygen (1 L·min^-1^) throughout the surgery. Rats were administered sub-cutaneous Meloxicam (2 mg·kg^-1^; Norbrook Inc., Lenexa, KA, United States) and Atropine (0.12 mg·kg^-1^; Westward Pharmaceutical co., Tujunga, CA United States) to provide long-term moderate analgesia and to prevent pleural effusion, respectively. Post-operatively rats are given Buprenorphine (0.1 mg·kg^-1^; Reckitt Benckiser, Slough, United Kingdom) to provide additional short-term analgesia. Rats were placed prone in a stereotaxic apparatus (Kopf Instruments, Tujunga, CA, United States) thermocoupled to a heating pad (TCAT 2-LV; Physitemp, Clifton, NJ, United States) and body temperature was maintained at a minimum of 33°C. Anaesthetic level was monitored throughout the surgery.

### EEG/EMG

For all animals, four electroencephalographic (EEG) electrodes were inserted into the brain, two cross-cortical electrodes which predominantly measure delta waves (**Figure 1A**; electrodes 1 and 2), and 2 cross-hippocampus electrodes which predominantly measure theta waves (**Figure 1A**; electrodes 3 and 4). Two electromyographic (EMG) wires were inserted into the trapezius muscle to record REM induced paralysis. The EEG electrodes (M1.4 x 3mm Philips Pan Head Machine Screws (DIN 7985H) – Stainless Steel (A2), SIP – M1.4-3-A2; Accu, Huddersfield, United Kingdom) and EMG wire are connected to a custom-built head mount fixed in place with Superbond dental cement (Prestige Dental, Bedworth, United Kingdom) and Vertex Orthoplast cold-curing orthodontic acrylic resin (Prestige Dental, Bedworth, United Kingdom). Animals were allowed to recover 2 weeks post-operative before recordings began, with food and water *ad libitum*, at ambient room temperature, 22 ± 2°C. Rats were pseudorandomised before every phase of testing and following the surgery the experimenters were blinded to the condition of each rat.

**Figure 1.**
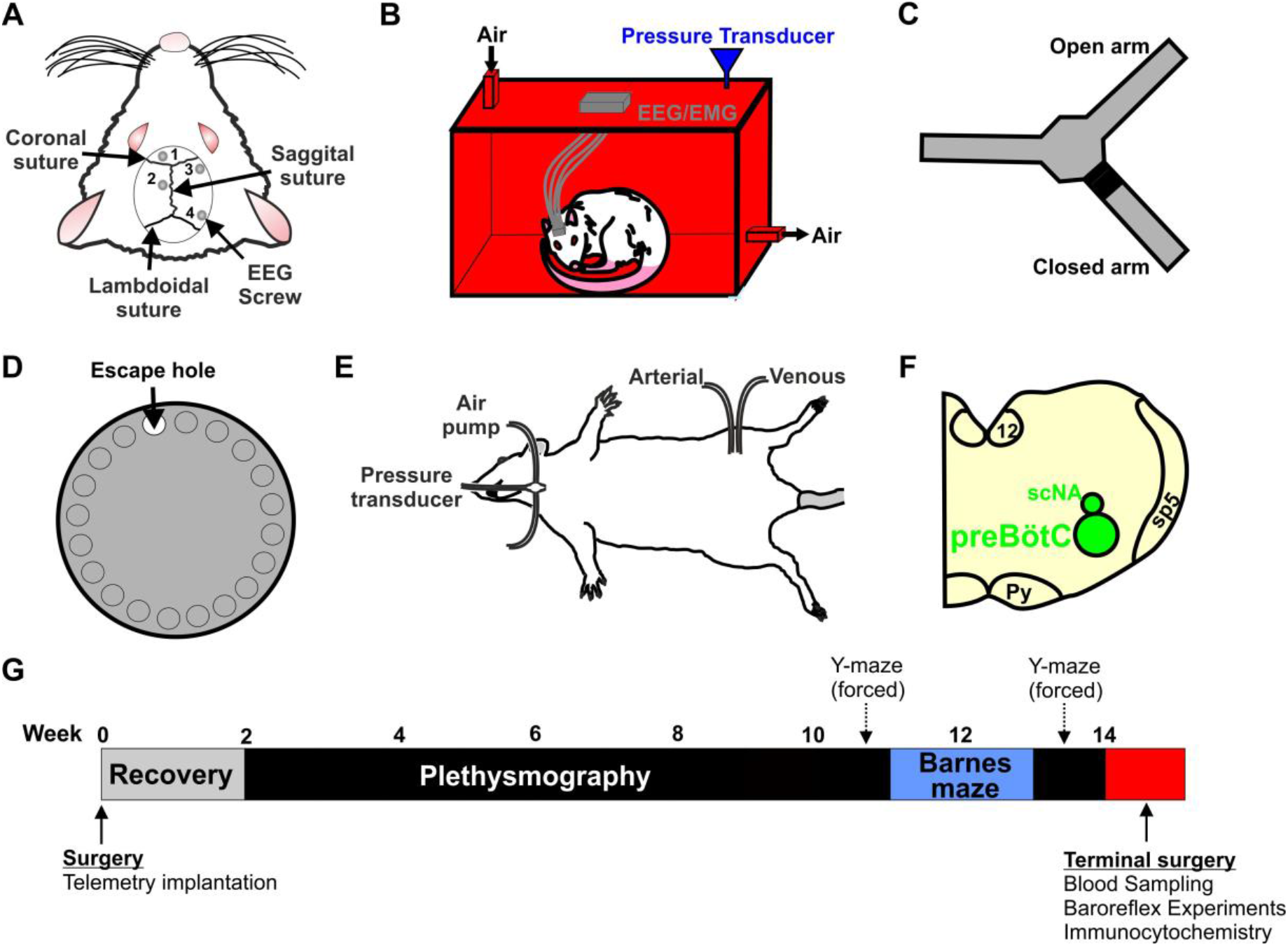
Testing innate sleep apnoea (SA) in SHR. **(A)** Electroencephalographic (EEG) and Electromyographic (EMG) electrodes were used to measure sleep-wake states in Spontaneously Hypertensive Rats (SHR) and Wistar Kyoto Rats (WKY). **(B)** A custom-built 4.5L plethysmograph was used to house the rats during the 3 h recording where EEG/EMG signals and flow (via a pressure transducer) were recorded. **(C)** Short-term memory was measured using a Y-Maze (forced alteration). **(D)** Long-term memory was measured using a Barnes Maze over a 12-day protocol. **(E)** Blood pressure, heart rate and breathing frequency recordings were taken whilst the rat was maintained under terminal anaesthesia and oxygen. **(F)** Immunocytochemical staining of the pre-defined area of the preBötzinger Complex (preBötC) and semi-compact Nucleus Ambiguus (scNA) was performed at the end of the experimental timeline. **(G)** Experimental timeline.

### Plethysmography

Rats were placed in to a 4.5 L plethysmography chamber (**Figure 1B**) with airflow set at 2 L·min^-1^ and calibrated via a 1 mL syringe, with an average ambient chamber temperature of 23 ± 0.9°C: the rats showed no signs of thermoregulation (panting, shivering, excessive locomotion, abnormal sleeping patterns). During the second week post-surgery, rats were allowed two acclimation sessions; once the plethysmograph was placed inside the rat’s home cage for 30 min, and on a separate occasion rats were placed into the plethysmograph for 30 min under experimental conditions. After these acclimation sessions, each rat underwent one 3 h recording per week for 7 weeks (postsurgical weeks 3-9 inclusive) during the light phase of the 12 h Light/dark cycle.

Airflow was recorded via a pressure transducer connected to the plethysmography chamber. Tidal volume (V_T_) was measured from trough-to-peak of each breath. Frequency (*f*) of breathing is measured from peak-peak of inspiration and is given as breaths·min^-1^. Minute ventilation (Ve) is determined by multiplying V_T_ x *f*. The respiratory parameters were measured during quiet wakefulness at the beginning of the recording. The signals from the pressure transducer were filtered and amplified using the Neurolog system via a 1401 interface (Digitimer, Welwyn Garden City, United Kingdom), and all data was acquired with Spike2 software (Cambridge Electronic Design, Cambridge, United Kingdom).

For assessment of the severity of SA, we measured the length of the respiratory disturbances (apnoea-hypoxia Index (AHI: sum of apnoeic + hypopnoeic events per hour of sleep)) by hand. Apnoea is defined as decrease in minute ventilation of 90% or greater for 1.8 s or more, hypopnoea is defined as decrease in minute ventilation of 50-90% for 1.8 s or more. Sighs (a compound breath followed by a period of apnoea) per hour of quiet wake + sleep were determined using EMG and flow traces; when the EMG was low with little/no movement shown combined with a clean period on the breathing trace this was considered quiet wake or sleep.

### Sleep-Wake Recordings

Theta waves were recorded from electrodes 1 (placed in the dorsal hippocampus) and 2 (placed in the frontal cortex) (**Figure 1A**), and delta waves were recorded from electrodes 3 and 4 (**Figure 1A**) then the EEG signals were amplified 7,500-fold and bandpass filtered at 5-70 Hz. EMG signals were recorded through the wire in the trapezoid muscle and amplified 4,500-fold and bandpass filtered at 50-1,500 Hz. All data was processed with the Spike2 software using the OSD script, and sleep scoring was aided with video footage from an HD camera of the rat throughout the entire 3 hour recording where physiological factors such as locomotion (WAKE), or paralysis (REM) are used to aid with analysis of sleep-wake state. EEG signals that had been band-pass filtered with a power spectrum between 0-4 Hz were classed as delta waves, and signals between 6-10 Hz were classed as theta waves. All EEG signals were smoothed with a 5 s time constant after being band-pass filtered. The EMG signals were then processed with a 5 s RMS constant, and the whole 3 h recording is separated into 5 s epochs.

Combining EEG and EMG analysis allowed us to determine sleep wake state into 4 distinct categories:

**WAKE**: low amplitude desynchronised EEG, high amplitude and frequency on EMG

**NREM**: high amplitude EEG delta waves, low amplitude and frequency on EMG

**REM**: low amplitude EEG theta waves and high theta:delta (T:D) ratio, low amplitude (or absent) EMG

**DOUBT**: any epoch that cannot be defined into one of the aforementioned states

All sleep state and AHI count analysis are expressed as per hour of sleep for consistency across the recordings, and sigh counts are expressed as per combined hour of sleep and quiet wakefulness.

### Cognitive Tests

#### Y Maze (forced alternation)

The Y maze (**Figure 1C**) was adapted from a radial arm maze displayed so that 3 arms (50 X 10 cm with 13 cm high walls) in the shape of the letter Y remained open (Stoelting, Dublin, Ireland). This phase of behavioural testing takes place either before or after the Barne’s maze in post-surgical week 10 or 13, the determination of which is pseudorandomised at the start of the experimental paradigm. The rat is placed into the maze in the entry arm, and either the left or the right arm is open, with the remaining arm closed. The location of the closed arm is pseudorandomised before each test. Spatial cues are placed at the end of each arm in plain sight of the rat, with comfortable ambient lighting (150-250 lux). Rats were given 10 mins to explore the maze, rested for a period of 1 h before being retested for 5 mins with both arms open. The order of the rats is pseudorandomised before the first test but is consistent between the tests. All test sessions are recorded with a camera (Henelec Model 335 BWL; Sony, Surrey, United Kingdom) that connects to a computer for offline analysis (Any-MAZE v4.96, Stoelting, Dublin, Ireland). Total time spent in each arm, total distance travelled in each arm and the total number of entries into each arm (defined as 20% of the rat’s whole body in the arm) were measured.

#### Barnes Maze

The Barnes maze is a circular maze (122 cm in diameter) with 19 false holes and one escape hole (9 cm in diameter; Stoelting, Dublin, Ireland) (**Figure 1D**). The maze is one metre off the ground. Spatial cues of different colours and shapes are spaced evenly around the maze at regular intervals in plain sight of the rat. Spatial clues are never placed directly over the escape hole. The surface of the table is brightly lit (>1500 lux) to create an adverse environment. On post-surgical week 10, rats underwent a 12-day Barnes maze protocol: Days 1-3 (learning phase) the escape hole contains an incentive (peanut butter); days 4-12 (acquisition phase) the incentive is provided in the home cage after the test, allowing the rat to be rewarded for completing the test but removing the olfactory stimulus from the maze. Before each test the order of the rats is pseudorandomised. The maze is thoroughly cleaned (70% ethanol) between tests to remove any traces of scent from the previous rat. No respiratory or other behavioural experiment occur on the days where the Barnes maze is performed to remove the impact these may have on the cognitive testing. The experiment is recorded by a camera (Henelec Model 335 BWL; Sony, Surrey, United Kingdom) connected to a computer for offline analysis (Any-MAZE v4.96, Stoelting, Dublin, Ireland). Exit errors (defined as the total number of times the rat investigated the escape hole but did not exit through it), total distance travelled on the maze and total time spent on the maze is measured. Search strategy was divided into 3 specific categories; sequential (travelling around the outside of the maze visiting ≥ 3 holes), spatial (travelling directly to within 2 holes of the escape hole and not visiting any other hole outside of this quadrant) or random (no specific pattern but must cross the centre line ≥ 2 times) [27].

### Baroreflex Sensitivity Test

Rats were anaesthetised in a gas inhalation chamber using 4% isoflurane (Piramal Healthcare, Mumbai, India) in pure oxygen and maintained throughout surgery with urethane (1 g/kg; Sigma: St. Louis, Missouri, United States) and α-chloralose (50 mg·kg^-1^ ;Sigma: St. Louis, Missouri, United States) diluted in standard sterile saline (0.9% NaCl). Additional doses of α-chloralose were administered as required. Anaesthetic levels are checked throughout with pedal withdrawal, and body temperature was maintained at 36.5°C by a thermocoupled heat mat. The trachea was cannulated. The femoral blood vessels were catheterised: the femoral vein was used for delivery of anaesthetics and the femoral artery was connected to pressure transducers (Digitimer, Welwyn Garden City, United Kingdom) to record blood pressure.

Baseline parameters were recorded for 20 mins. Phenylephrine (3.5 mg·mL^-1^; Fisher Scientific UK, Loughborough, Leicestershire, LE11 5Rg, United Kingdom) or sodium nitroprusside (35 mg·mL^-1^; Honeywell, Chem-Supply Pty Ltd, 38-50 Bedford Street, Gillman SA 5013, Australia) were administered followed by a saline flush. The animal was allowed to rest until measurements returned to baseline, or a new baseline was established over 20 mins before administration of the second drug. The order of the drugs was pseudorandomised before the surgery starts. Mean arterial pressure (MAP) was calculated as [⅔ Diastolic pressure + ⅓ Systolic Pressure] and heart rate (BPM) was calculated as peak to peak of blood pressure and is given as beats·min^-1^. Data are displayed as the ratios of the MAP or heart rate after injection compared to the average baseline of each animal.

### Tissue Collection

At the end of the baroreflex-sensitivity tests rats were transcardially perfused with 4% paraformaldehyde (PFA). The cortex and the medulla were removed and fixed overnight in 4% PFA at 4°C. Once the brains had sunk (typically 2-3 days) they were transferred to, and stored in, cryoprotectant (30% sucrose + 0.02% sodium azide) at 4°C.

### Immunocytochemistry

Tissue was sectioned to 50 μm on a cryostat (Bright instruments Ltd, Huntington, United Kingdom). Slices were washed in PBS for six x 5-min washes, before being placed in a sodium citrate buffer (tri-sodium citrate (dihydrate) 2.94g, 1 L ddH_2_O, 0.5 mL Tween 20, adjusted to pH 9.0 with NaOH), preheated to 80°C in a still water bath for (heat-induced) antigen retrieval. The free-floating slices were left for 30 minutes, before being washed in PBS for 6 x 5-minute washes. Tissue was transferred to blocking solution (PBS containing 0.1% Triton X and 5% bovine-serum albumin) to block non-specific binding of antibodies for 1 h at room temperature. The slices were then incubated overnight at room temperature in the blocking solution plus the following primary antibodies:

#### Medulla

preBötC counts: mouse anti-NeuN (1:100, MAB337, Merck Millipore, Watford, United Kingdom), rabbit anti-NK1R (1:500, ab5060, Merck Millipore, Watford, United Kingdom).

Microvascular rarefaction: rabbit anti-VCAM (1:200, ab134047, Abcam, Cambridge, United Kingdom), mouse anti-Tie2/TEK (1:200, ab33, Merck Millipore, Watford, United Kingdom), goat anti-ChAT (1:50, ab144P, Merck Millipore, Watford, United Kingdom).

#### Hippocampus

##### Neuroinflammation

goat anti-Iba-1 (1:83, ab5076, Abcam, Cambridge, United Kingdom).

The slices were then washed in PBS for six x 5-minute washes, then placed in blocking solution for 1 h at room temperature. The slices were then incubated for 2 h at room temperature with the following secondary antibodies:

#### Medulla

preBötC counts: donkey anti-mouse Alexa Fluor 568 (1:250, A10037, Invitrogen, Waltham, MA, United States), donkey anti-rabbit Alexa Fluor 488 (1:250, A10037, Invitrogen, Waltham, MA, United States).

Microvascular rarefaction: donkey anti-rabbit Alexa Fluor 568 (1:250, ab150074, Abcam, Cambridge, United Kingdom), donkey anti-mouse Alexa Fluor 488 (1:250, A-21202,Hampton, Fisher Scientific, New Hampshire, United States), donkey anti-goat Alexa Fluor 405 (1:250, ab175665, Abcam, Cambridge, United Kingdom);

#### Hippocampus

##### Neuroinflammation

donkey anti-goat Alexa Fluor 488 (1:250, A10037, Invitrogen, Waltham, MA, United States), DAPI staining solution (1:1000, ab228549, Abcam, Cambridge, United Kingdom).

All slices were then mounted on polylysine microscope slides and dehydrated overnight at room temperature. The slides were then rehydrated in ddH_2_O, mounted (Cytoseal 60, Electron Microscopy Sciences, Hatfield, PA, United States) and coverslipped.

Slides were imaged on a confocal microscope (Zeiss 880, Zeiss, Jena, Germany) using Zen Blue and Zen Black software (Zeiss, Jena, Germany).

#### Medulla analysis

The number of NeuN-positive cells were counted to determine viral transduction efficacy over a 600 μm diameter circle below the semi-compact nucleus ambiguous [28, 29], defining the preBötC. Following a background subtraction on the staining for either NK1R-positive, VCAM-positive or Tie2/TEK-positive neurons, fluorescence intensity was measured within the preBötC. The same intensity analysis and cell count analysis was performed on the nucleus ambiguus for the NK1R and NeuN analysis to confirm the continuity of staining across all tissue.

#### Hippocampus analysis

the dentate gyrus or the CA1 region were imaged. To identify activated microglia, we counted DAPI-stained nuclei encapsulated with Iba1 anywhere in the visual field. Given we only stained every second slice, nuclei could only be present in a single section, and thus removing overestimation by double counting.

All image processing and analysis was performed on ImageJ (ImageJ, U.S. National Institutes of Health, Bethesda, MA, United States).

### Experimental Design

For each stage of the experimental timeline, the order of the rats was pseudorandomised, and for the duration of the experimental timeline, the experimenter(s) were blinded to the condition (hypertensive vs normotensive). The plethysmograph acclimation occurred in the 2^nd^ week post-surgery, and the experimental plethysmography recordings started in the 3^rd^ week post-surgery and took place weekly up to and inclusive of the 9^th^ week post-surgery. Rats then either underwent a 1-day Y maze protocol or a 12-day Barnes maze protocol, the order of which was pseudorandomised for each batch of rats. The other maze was then completed after a 3-day rest period. One week after the final maze session, the baroreflex sensitivity experiments were performed, and tissue was collected, and. Once all experiments were finished and all analysis was completed, then the experimenters were unblinded (**Figure 1G**).

### Data analysis

All experimental units are a rat, and all technical repeats are averaged to create a biological repeat for analysis. An Iglewicz & Hoaglin’s robust test (with a modified Z score of ≥ 3.5 https://contchart.com/outliers.aspx) was used to remove outliers before statistical analyses were carried out. To establish the state of normality, Shapiro-Wilks tests were carried out on the sham-operated groups. All experiments are recorded and analysed using Spike software (vs7.08, Cambridge Electronic Design).

Data for baseline MAP (*p* = 0.9), Pe MAP (*p* = 0.3), SNP MAP (*p* = 0.2), baseline HR (*p* = 0.2), Pe HR (*p* = 0.8), SNP HR (*p* = 0.6), preBötC cell count (*p* = 0.5), Tie2/TEK preBötC staining intensity (*p* = 0.8), VCAM1 preBötC staining intensity (*p* = 0.1), BötC staining intensity (*p* = 0.2), BötC cell count (*p* = 0.9), NA NK1R staining intensity (*p* = 0.9), NA cell count (*p* = 0.9), V_T_ (*p* = 0.7), V_E_ (*p* = 0.1), *f* (*p* = 0.9), average sighs (*p* = 0.2), number of dyspnoeic episodes during sleep (*p* = 0.9), time spent dyspnoeic (*p* = 0.3), time spent in disordered breathing by sleep state (WKY: *p* = 0.9; SHR: *p* = 0.9), Wake (*p* = 0.8), NREM (*p* = 0.5), REM (*p* = 0.8) were deemed Gaussian by a Shapiro-Wilks test for normality and tested by a two sample t test. Data are expressed as mean ± SD.

Data for preBötC NK1R staining intensity (*p* = 0.003), CA1 Iba1 positive cell count (*p* = 0.05), DG Iba1 positive cell count (*p* = 0.00007), Barnes maze search strategies (*p* = 0.003) were deemed non-Gaussian by a Shapiro-Wilks test for normality and tested by a Kruskal Wallis ANOVA with Dunn-Sidak post-hoc correction. Data are expressed as mean ± SD.

Data for average length of dyspnoeic episode (*p* = 0.4) was deemed Gaussian by a Shapiro-Wilks test for normality and tested by a two-way ANOVA with Bonferroni correction. Data are expressed as mean ± SD.

Data for Y Maze entries (*p* = 0.008) was deemed non-Gaussian by a Shapiro-Wilks test for normality and tested by a two-way repeated measures ANOVA with Bonferroni correction. Data are expressed as mean ± SD.

Data for number of dyspnoeic episodes by week (*p* = 0.3) and time spent dyspnoeic by week (*p* = 0.5), Y Maze duration (*p* = 0.9), Y Maze distance (*p* = 0.8), Y Maze discrimination ratio (distance (*p* = 0.5), duration (*p* = 0.2), entries (*p* = 0.2)), were deemed Gaussian by a Shapiro-Wilks test for normality and tested by a two-way repeated measures ANOVA with Bonferroni correction. Data are expressed as mean ± SD.

Data for Barnes maze duration (*p* = 0.4) was deemed non-Gaussian, Barnes maze distance (*p* = 0.1), and Barnes maze exit errors (*p* = 0.1) were deemed Gaussian by a Shapiro-Wilks test for normality and tested by a two-way repeated measures ANOVA with a Dunn-Sidak correction. Data are expressed as mean ± SD.

## Results

### SHRs are innately hypertensive compared to WKY

To confirm SHRs displayed a hypertensive phenotype, we performed blood pressure recordings. SHRs had increased blood pressure (WKY: 97 ± 14.1 MAP, n = 4 vs SHR: 188.2 ± 16.1 MAP, n = 3, *p* = 0.001, **Figure 2A**), but there was no difference in heart rate between the groups (WKY: 128.8 ± 18.4 BPM, n = 3 vs SHR: 176.1 ± 66.3 BPM, n = 5, **Figure 2B**). Therefore, hypertension in our SHRs is uncompensated, indicative that the animals have been hypertensive for a long period of time [30],

**Figure 2.**
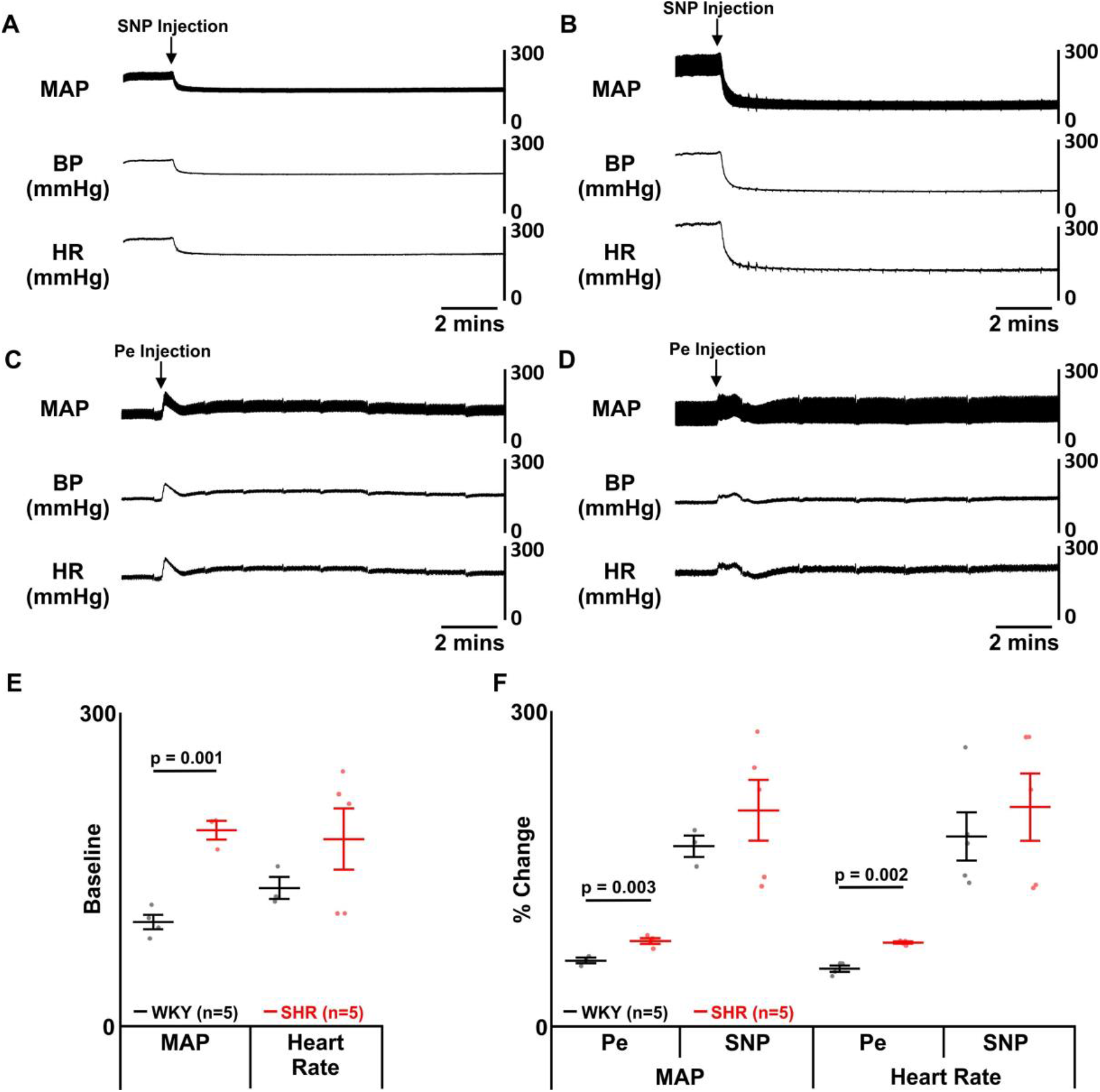
SHRs *have higher blood pressure and altered baroreflex sensitivity*. **(A)** Representative trace showing mean arterial pressure (MAP), blood pressure (BP) and heart rate (HR) in a Wistar Kyoto (WKY) rat after injection of sodium nitroprusside (SNP) **(B)** Representative trace showing mean arterial pressure (MAP), blood pressure (BP) and heart rate (HR) in a Wistar Kyoto (WKY) rat after injection of phenylephrine (Pe) **(C)** Representative trace showing mean arterial pressure (MAP), blood pressure (BP) and heart rate (HR) in a spontaneously hypertensive (SHR) rat after injection of sodium nitroprusside (SNP) **(D)** Representative trace showing mean arterial pressure (MAP), blood pressure (BP) and heart rate (HR) in a spontaneously hypertensive (SHR) rat after injection of phenylephrine (Pe). **(E)** Group data showing blood pressure and heart rate before injections (baseline) in SHRs (red) and WKY rats (black). **(F)** Group data showing blood pressure and heart rate after injections of phenylephrine (Pe) and sodium nitroprusside (SNP) in SHRs (red) and WKY rats (black). Data are represented as mean ±SD with individual data points.

We then tested if SHRs displayed altered baroreflex sensitivity. SHRs showed an increase in baroreflex sensitivity after phenylephrine (Pe) injection for both blood pressure (Pe WKY: 57.4 ± 6.2 MAP, n = 4 vs SHR: 82.8 ± 2.3 MAP, n = 3, *p* = 0.002, **Figure 2A**) and heart rate (Pe WKY: 59.9 ± 4.8 BPM, n = 3 vs SHR: 82.3 ± 5.6 BPM, n = 4, *p* = 0.003, **Figure 2B**). Therefore, there is an increased sensitivity to vasoconstriction. We did not see a difference between groups after injection of sodium nitroprusside (SNP) for either blood pressure (SNP WKY: 186.6 ± 52.8 MAP, n = 5 vs SHR: 215.3 ± 73.7 MAP, n = 5, **Figure 2A**) or heart rate (SNP WKY: 199.9 ± 54.4 BPM, n = 4 vs SHR: 208.2 ± 65.8 BPM, n = 5, **Figure 2B**). Therefore, SHRs display weakened arterial walls as a result of prolonged hypertension.

### Innate SA leads to sleep disordered breathing without alterations in respiration during wakefulness

To determine if SHRs displayed innate sleep apnoea, we measured the breathing of WKY and SHRs with plethysmography (**Figures 1B, 3A-J**) and sleep-wake state via EEG/EMG recordings (**Figures 1A, 3A**). AHI (sum of apnoeic + hypopnoeic events per hour of sleep) was increased in SHRs (WKY: 9 ± 0.6 incidences·hour of sleep^-1^, n = 9 vs SHR: 20.7 ± 1.1 incidences·hour of sleep^-1^, n = 6; p = 0.00000005; **Figure 3B**) as was total duration of apnoeas + hypopnoeas (WKY: 22.4 ± 0.8 s, n = 9 SHR: 44.5 ± 2.2 s, n = 8; p = 0.00000005; **Figure 3B**). Sleep apnoea in WKY rats was no different to Sprague Dawley (SD) controls from [29], for AHI (SD: 9.4 ± 0.8 incidences·hour of sleep^-1^, n = 14 vs WKY: 9.2 ± 0.6 incidences·hour of sleep^-1^, n = 9; **Figure 3C**) or total time spent dyspnoeic (SD: 26.2 ± 1.5 s, n = 14 WKY: 22.9 ± 0.8 s, n = 9; **Figure 3C**).

**Figure 3.**
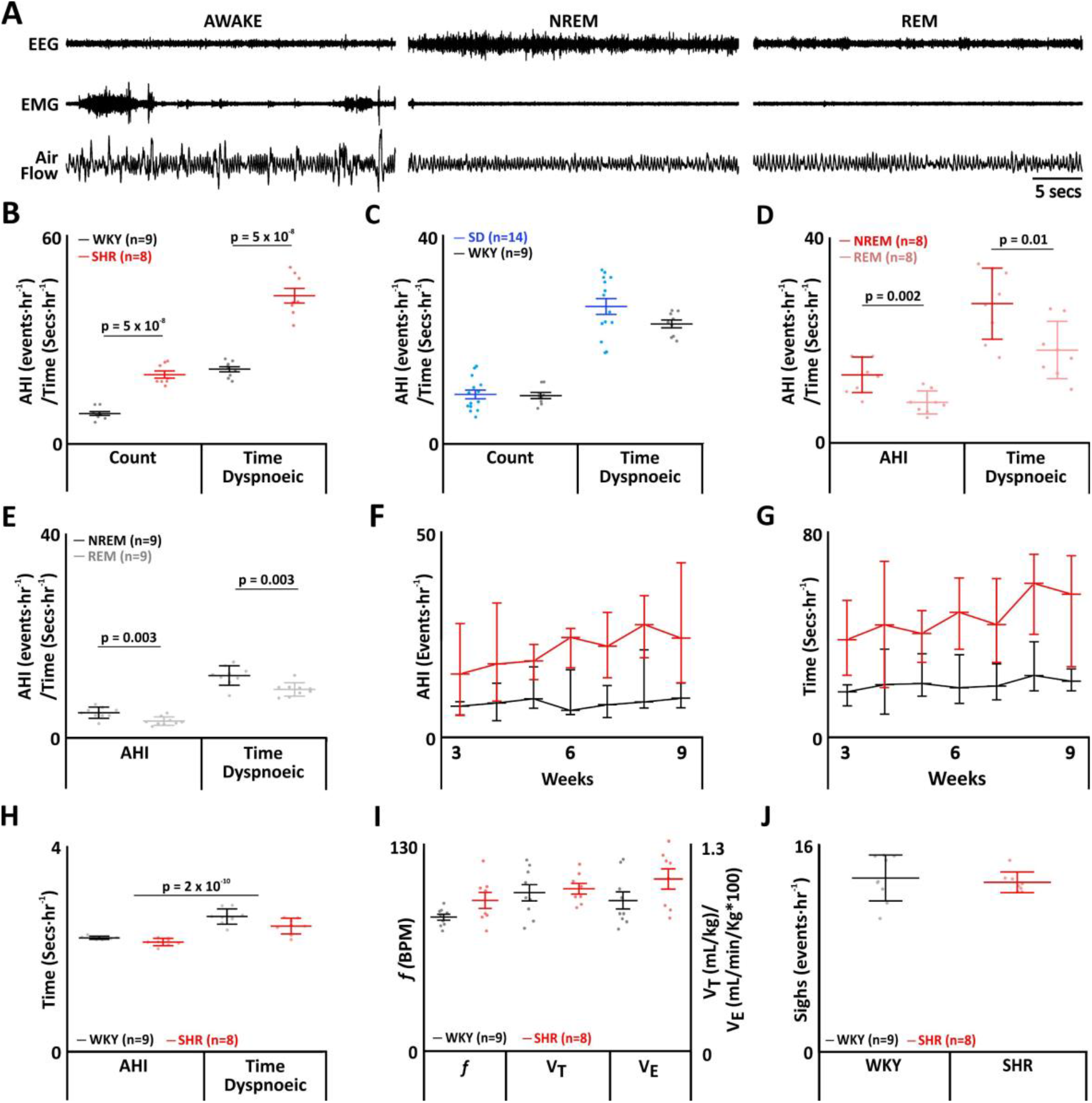
SHR phenotype displays apnoeas with no change in breathing. **(A)** Plethysmograph traces from SHRs, showing respiratory movements as measured by changes in air flow in a plethysmograph, with EEG and EMG electrodes for assigning sleep-wake state. **(B-J)** Data represents WKY rats (black) compared to SHRs (red). **(B)** Frequency of breathing disturbances (apnoea-hypopnoea index (AHI)) and duration spent dyspnoeic (time) between WKYs and SHRs. **(C)** Frequency of breathing disturbances (apnoea-hypopnoea index (AHI)) and duration spent dyspnoeic (time) between SDs and WKYs. **(D)** Frequency of breathing disturbances and duration spent dyspnoeic by sleep state, non-rapid eye movement (NREM) (red) and rapid eye movement (REM) (pink), in SHRs. **(E)** Frequency of breathing disturbances and duration spent dyspnoeic by sleep state, NREM (black) and REM (grey) in WKY rats. **(F)** Progression of AHI by week **(G)** Progression of time spent dyspnoeic by week. **(H)** . Average duration of a dyspnoea during NREM and REM. **(I)** Respiratory parameters during wakefulness. Left axis shows values for frequency (f), right axis for tidal volume (V_T_) and minute ventilation (V_e_). **(J)** Frequency of sighing. Data are represented as mean ±SD with individual data points.

For both WKYs and SHRs, apnoeas occurred more in NREM sleep than REM sleep (WKY: NREM: 5.4 ± 0.4 incidences·hour of sleep^-1^, n = 9 vs REM: 3.8 ± 0.4 incidences·hour of sleep^-1^, n = 9; p = 0.003; SHR: NREM: 12.8 ± 1.1 incidences·hour of sleep^-1^, n = 8 vs REM: 7.9 ± 0.8 incidences·hour of sleep^-1^, n = 8; p = 0.002; **Figure 3D,E**). There was also an increase in time spent dyspnoeic during NREM (WKY: NREM: 12.7 ± 2 s, n = 9 vs REM: 9.9 ± 0.4 s, n = 9, p = 0.003; SHR: NREM: 26.3 ± 2.3 s, n = 8 vs REM: 18.2 ± 2 s, n = 8; p = 0.01; **Figure 3D,E**).

As the weeks progressed there was no difference seen in the number of dyspnoeic episodes or duration of the dyspnoeic events (**Figure 3F,G**), and there was no difference seen between the average duration of dyspnoeic events between groups (WKY: 2.4 ± 0.2 s, n = 9 vs SHR: 2.3 ± 0.2 s, n = 8; **Figure H**).

There was no difference in respiratory parameters during wakefulness between the groups, both maintained *f* (WKY: 87 ± 15 breaths·min^-1^, n = 9 vs SHR: 98 ± 6 breaths·min^-1^, n = 8; **Figure 3I**), V_T_ (WKY: 1.0 ± 0.1 mL·kg^-1^, n = 9 vs SHR: 1.0 ± 0.0 mL·kg^-1^, n = 9; **Figure 3I**), and V_e_ (WKY: 89 ± 5 mL·kg^-1^·min^-1^, n = 9 vs SHR: 101 ± 6 mL·kg^-1^·min^-1^, n = 8; **Figure 3I**). There was also no difference in the sigh rate between groups (WKY: 13 ± 1.8 incidences·h^-1^, n = 9 vs SHR: 13 ± 0.8 incidences·h^-1^, n = 8; **Figure 3J**), showing overall integrity of the respiratory microcircuit remained intact.

### SHRs have reduction in neurons in the preBötC

Previously, we determined that a 23% loss of neurons in the preBötC led to the development of moderate SA [29]. To assess if the sleep apnoea seen in SHRs is due to an impairment of the preBötC, we conducted immunocytochemical analysis of the preBötC and NA. There was a loss of presumptive rhythm generating cells in the preBötC in SHRs (WKY: 29.6 ± 7.0 NK1R intensity, n = 8 vs SHR: 16.5 ± 4.1 NK1R intensity, n = 8; p = 0.002; **Figures 4A,G**) with no overall difference in intensity in the NA stained cells (WKY: 37.5 ± 4.3 NK1R intensity, n = 8 vs SHR: 35.0 ± 4.8 NK1R intensity, n = 8; **Figures 4A,G**), nor BötC cells (WKY: 18.0 ± 0.9 NK1R intensity, n = 7 vs SHR: 17.4 ± 0.7 NK1R intensity, n = 6; **Figures 4C,G**). We also saw a loss of neurons in SHRs (WKY: 1745 ± 179 neurons, n = 8 vs SHR: 1164 ± 143 neurons, n = 7; p = 0.000001; **Figures 4B,I**) with no overall loss of NA neurons (WKY: 202 ± 37 neurons, n = 8 vs SHR: 199 ± 35 neurons, n = 7; **Figures 4B,I**) nor BötC neurons (WKY: 1374 ± 105 neurons, n = 7 vs SHR: 1547 ± 148 neurons, n = 6; **Figures 4D,I)**. A loss of 33% of preBötC exceeds the threshold required to induce sleep apnoea and explains why SHRs are afflicted by it.

**Figure 4.**
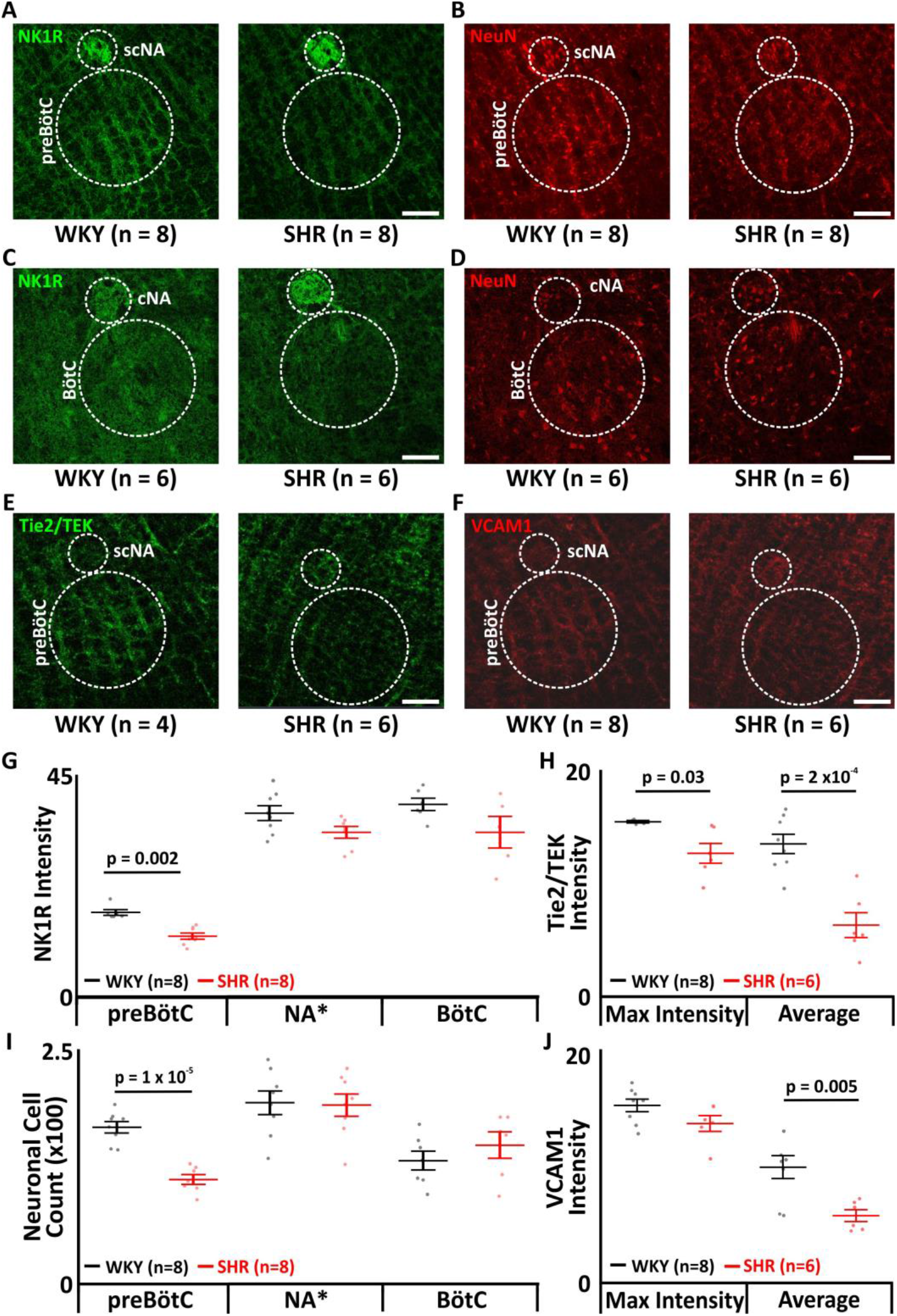
SHR phenotype displays a neuronal and vascular deficit in the pre-defined area of the preBötC. **(A)** Micrographs showing immunocytochemical staining of neurokinin 1 receptor (NK1R) positive cells (NK1R: green) of the preBötzinger Complex (preBötC) and semi-compact nucleus ambiguus (scNA). **(B)** Micrographs showing immunocytochemical staining of the neurones (NeuN: red) of the preBötzinger complex (preBötC) and semi-compact nucleus ambiguus (scNA). **(C)** Micrographs showing immunocytochemical staining of neurokinin 1 receptor (NK1R) positive cells (NK1R: green) of the Bötzinger Complex (BötC) and compact nucleus ambiguus (cNA). **(D)** Micrographs showing immunocytochemical staining of the neurones (NeuN: red) of the Bötzinger complex (BötC) and compact nucleus ambiguus (cNA). **(E)** Micrographs showing immunocytochemical staining of Tie2/TEK positive cells (Tie2/TEK: Green) of the preBötzinger Complex (preBötC) and semi-compact nucleus ambiguus (scNA). **(F)** Micrographs showing immunocytochemical staining of vascular cell adhesion molecule 1 (VCAM1) positive cells (VCAM1: Red) of the preBötzinger Complex (preBötC) and semi-compact nucleus ambiguus (scNA). **(G)** Group data of the NK1R stain intensities in the preBötC NA and BötC show in **(A)** and **(C). (H)** Group data of the maximum intensity Tie2/TEK preBötC staining intensity represented by the micrographs shown in **(E)** and average preBötC staining intensity (no micrograph shown). **(I)** Group data of the neuronal cell counts of the preBötC NA and BötC show in **(B)** and **(D). (J)** Group data of the maximum intensity VCAM1 preBötC staining intensity represented by the micrographs shown in **(F)** and average preBötC staining intensity (no micrograph shown). Data are represented as mean ±SD with individual data points.

To understand why neurons within the preBötC were dying, we investigated vascular health in the region. We found a loss of vasculature in the preBötC (WKY: 15.6 ± 0.5 Tie2/TEK intensity, n = 6 vs SHR: 12.6 ± 0.9 Tie2/TEK intensity, n = 6; p = 0.03; **Figures 4E,H**). We also detected a reduction in VCAM1 (WKY: 11.1 ± 0.9 VCAM1 intensity, n = 8 vs SHR: 7.2 ± 1.1 VCAM intensity, n = 6; p = 0.005; **Figures 4,J**), though maximal intensity remained unchanged (WKY: 15.9 ± 0.5 VCAM1 intensity, n = 8 vs SHR: 14.5 ± 0.6 VCAM intensity, n = 6; ns; **Figures 4F,J**). Given the vasculature showed the same level of damage (as seen through maximal intensity), the reduction in overall VCAM is due to the loss of vasculature (as seen with Tie2/TEK staining).

### SHR phenotype display no change in time spent in REM

Dyspnoeas were observed during both phases of sleep, and dyspnoeas induce microarousals, so we tested whether SHRs exhibited any altered or disturbed sleep patterns. There were no sleep disturbances seen in either group for either sleep state (WAKE: WKY: 49 ± 6%, n = 9 vs SHR: 52 ± 8%, n = 8; **Figure 5C**; NREM: WKY: 46 ± 6%, n = 9 vs SHR: 42 ± 7%, n = 8; **Figure 5C**; REM: WKY: 5 ± 1%, n = 9 vs SHR: 5 ± 1%, n = 8; **Figure 5C**).

**Figure 5.**
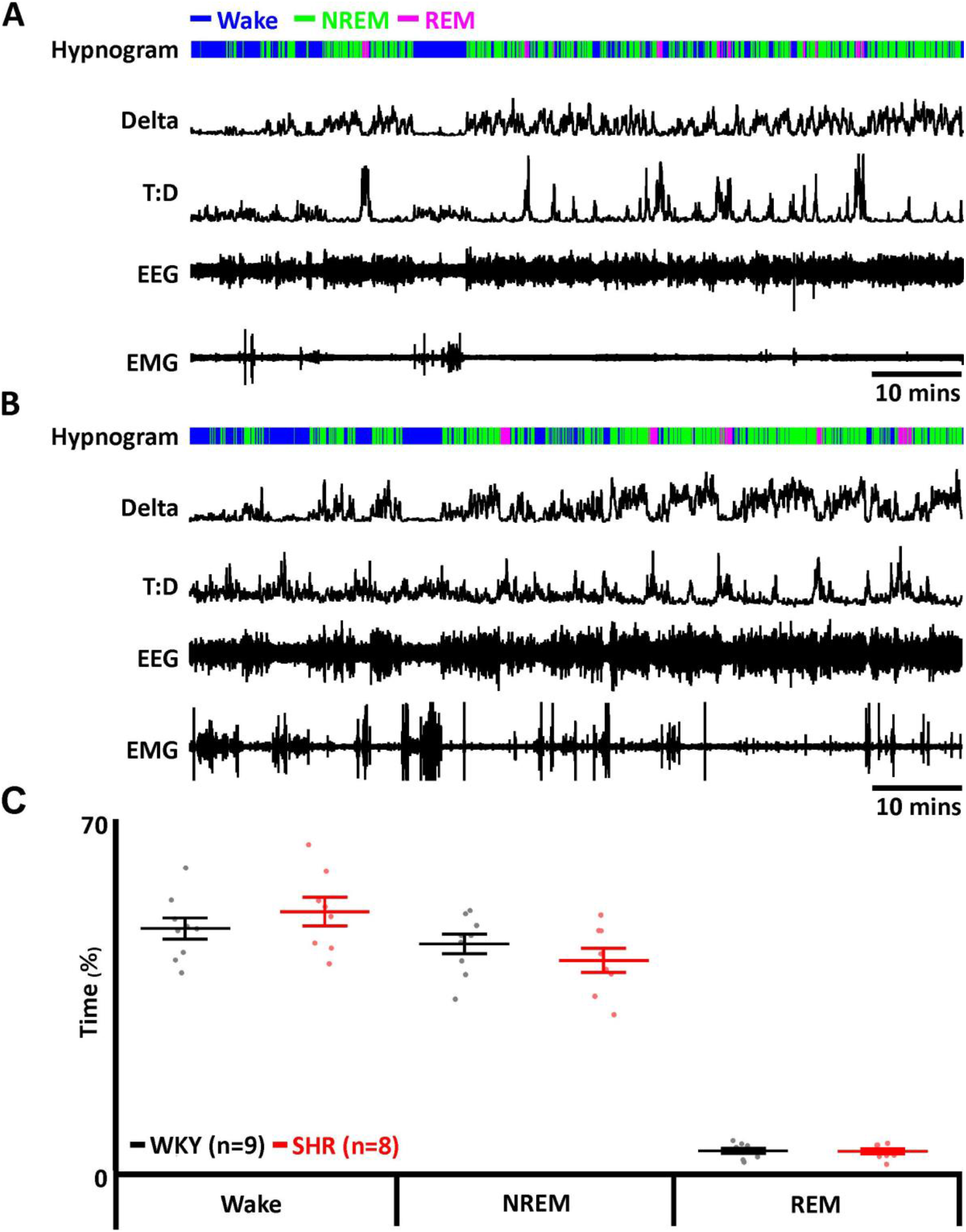
SHR phenotype display no change in time spent in REM. **(A)** Data from a representative Wistar Kyoto (WKY) rat. **(B)** Data from a representative spontaneously hypertensive (SHR) rat. **(A**,**B)** EMG recordings were used to determine activity and to identify periods of REM and NREM sleep. EEG recordings were used to determine delta wave activity for NREM sleep and to calculate theta-delta ratio (T:D) for REM sleep. Hypnograms show time spent awake, and in REM and NREM sleep stages. **(C)** Time spent in wakefulness, and NREM and REM sleep in SHRs (red) compared to WKY rats (black). Data are represented as mean ±SD with individual data points.

Whilst it is surprising therefore that we did not see altered sleep, sleep fragmentation in SHRs occurs in higher frequency at the end of the light phase [31]. Therefore, the lack of sleep fragmentation seen here is because we were only able to capture a snapshot of the animal’s sleep that did not always span this period.

### SHRs have neuroinflammation

Sleep apnoea induces neuroinflammation [29], thus we next investigated if SHRs showed elevated levels of activated microglial in the hippocampus. SHRs showed neuroinflammation through increased activated microglia in the hippocampus in both the CA1 region (WKY: 4 ± 6 cells, n = 6 vs SHR: 36 ± 16 cells, n = 9: p = 0.004; **Figures 6A,B**) and dentate gyrus (WKY: 5 ± 6 cells, n = 5 vs SHR: 40 ± 10 cells, n = 9: p = 0.004; **Figures 6C,D**).

**Figure 6.**
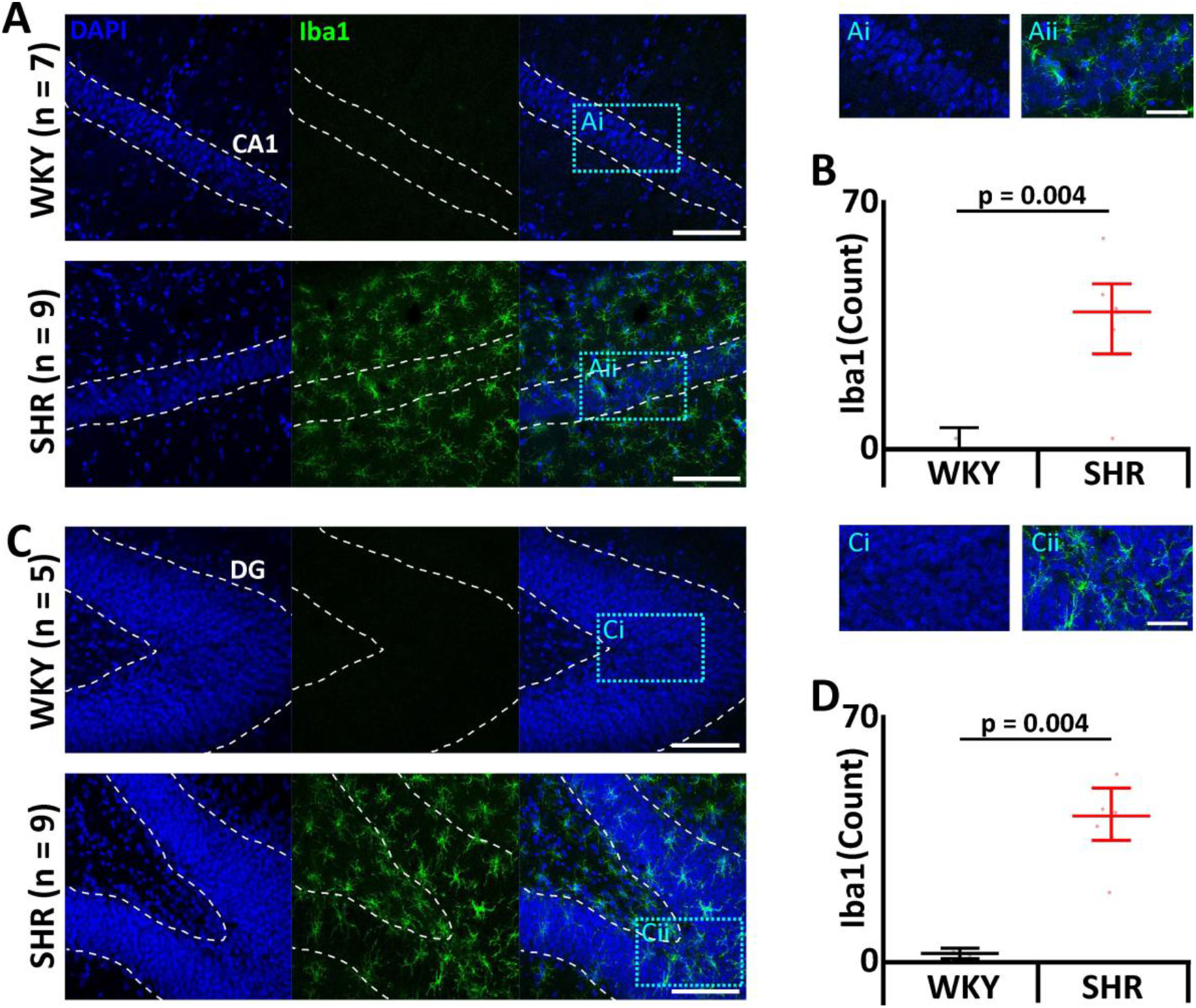
SHR phenotype displays elevated levels of activated microglia. **(A**,**C)** Micrographs showing activated microglia, a marker of neural inflammation, through ionized calcium binding adaptor molecule 1 (Iba1) expression. Scale bars 100μm. Blue boxes represent expanded areas shown in i and ii. Expanded micrograph scale bars 50μm. **(A)** CA1 region. **(B)** Dentate gyrus (DG). **(B**,**D)** Group data displayed in **(B)** CA1 region and **(D)** DG. Data are represented as mean ±SD with individual data points.

### SHRs have deficits in long term memory

We next investigated the effect of neuroinflammation on long-term memory via the Barnes maze. SHRs spent a similar amount of time on the maze (WKY: 39 ± 10 s vs SHR: 33 ± 11 s; mean difference 6; std error 15; DF = 14; **Figures 7A**) but cover more distance (WKY: 1.7 ± 0.1 m vs SHR: 2.5 ± 0.2 m; mean difference -8; std error 2.2; DF = 14; p=0.003, **Figures 7A**). and have more escape hole failures (WKY: 0.7 ± 0.1 vs SHR: 1.7 ± 0.1, mean difference -0.2; std error 0.2; DF = 14 p=0.00001; **Figures 7A**). Therefore, SHRs have altered long-term memory.

**Figure 7.**
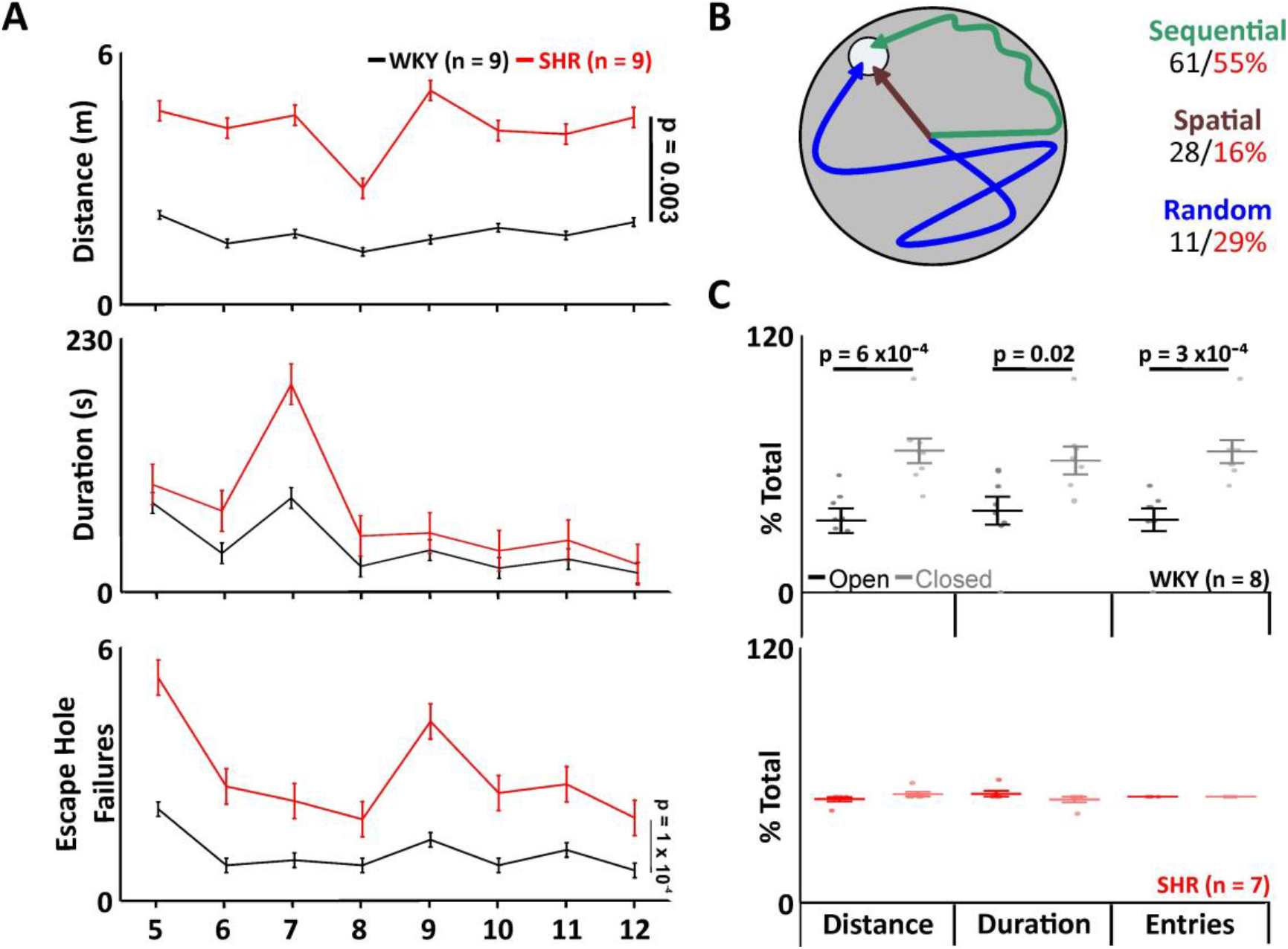
SHR phenotype displays short-term and long-term memory decline. **(A-B)** Barnes Maze. **(A)** Group data showing time spent and distance covered on the maze, as well as the number of escape hole failures in SHRs (red) compared to WKY rats (black). **(B)** Cartoon displays search strategies used by SHRs (red) compared to WKY rats (black). **(C)** Group data showing number of entries, distance covered, and time spent in the novel arm (light shading) compared to the previously open arm (dark shading) in SHRs (red) compared to WKY (black). Data are represented as mean ±SD with individual data points.

There was no difference in search strategies between WKY or SHR (Spatial -WKY: 28% ± 6% vs SHR: 16% ± 3%: sequential - WKY: 61% ± 8% vs SHR: 55% ± 6%: random - WKY: 11% ± 5% vs SHR: 29% ± 5%; **Figure 7B**).

### SHRs have deficits in short term memory

As neuroinflammation contributes to cognitive impairments through the depletion of synapses and their plasticity [32] we wanted to determine if this impacted learning and short-term memory of SHRs. In a Y-maze forced alteration test), WKYs, but not SHRs displayed a preference to the novel arm for distance travelled (WKY: Open: 34% ± 7%; Closed: 66% ± 7%; p = 0.0006; n = 8 and SHR: 52% ± 1%; Closed: 48% ± 1%; n = 7; **Figure 7C**), the duration spent in the novel arm (WKY: Open: 39% ± 7%; Closed: 61% ± 7%; p = 0.02; n = 8 and SHR: 54% ± 1%; Closed: 46% ± 1%; n = 7; **Figure 7C**) and entries into the novel arm (WKY: Open: 34% ± 6%; Closed: 66% ± 6%; p = 0.0003; n = 8 and SHR: 52% ± 0%; Closed: 48% ± 0%; n = 7; **Figure 7C**).

Both groups covered the same total distance in the maze, so the data collected was not affected by any locomotive or anxiety shown by the rats (WKY: 8.9 m ± 2.2 m, n = 6 vs. SHR: 9 m ± 2.2 m, n = 7, data not shown).

## Discussion

Hypertension is referred to as a ‘silent killer’, causing blood vessel damage and vascular remodelling, affecting all major organs including the brain. Over time, blood vessels become thicker, reducing elasticity and causing a reduction in the inner diameter of the vessel, therefore increasing vascular resistance [33]. This combination of atherosclerosis and hypertension leads to microvascular rarefaction [34], the loss of small blood vessels comprising microcirculation. Microvascular rarefaction has long been associated with symptoms of hypertension [35]. The reduction in microvascular density is caused either by reduced angiogenesis, which is seen more commonly in cases of genetic hypertension, or vessel destruction which is more commonly seen in cases of secondary hypertension. In the brain, this causes cerebral small vessel disease, which if left untreated can progress into cognitive decline [36].

Interestingly, the majority of the apnoeas we saw in NREM sleep were terminated by arousal, which is indicative of obstructive sleep apnoea [37]. The recurrent laryngeal nerve can be damaged as a result of hypertension via inflammation caused by increased levels of aldosterone causing oedema of the tissue in the upper and lower airways [38]. This nerve innervates the muscles of the larynx, the main role of which is to protect the lower respiratory tract to allow basic functions such as talking, swallowing and breathing [39]. Damage to this fundamental nerve leads to reduced muscle tone, allowing the tongue to fall to the back of the mouth during sleep, obstructing airflow resulting in obstructive sleep apnoea [40, 41]. Obstructive sleep apnoea is defined by a dyspnoeic episode where respiratory effort is still present, and the apnoeas are typically terminated by micro-arousals resulting in fragmented sleep, and a reduction in slow wave and REM sleep [42].

We found evidence of microvascular rarefaction in the preBötC in our model, as there was a reduced intensity of Tie2/TEK in SHRs. Tie2/TEK is a tyrosine kinase receptor central to vascular stability [43]. This receptor has three ligands (Ang1, Ang2 and Ang4), each angiopoietins, which contribute to the downstream signalling to complement the VEGF pathway by impacting vascular development [44, 45]. We postulate the density of microvasculature in the preBötC has been reduced through diminished angiogenesis over a prolonged period. VCAM1 upregulates the transmigration of leukocytes across membranes [46]. A loss of VCAM1 is indicative of overall loss of tissue, most likely due to inappropriate activation of the cell adhesion molecule enhancing chronic inflammation seen in that area [47]; in this case, the preBötC. Interestingly, whilst we saw a 39% decrease in NK1R positive neurons and 33% decrease in preBötC neurons overall, we saw no loss of neurons in the BötC, and no change in the number of sighs per hour of sleep, which suggests that the damage caused by microvascular rarefaction is differentially regulated throughout the respiratory column. Whether this selective response is regulated through differential sensitivity to oxidative stress, hypertension, or other factors remains unknown.

Inhibitory preBötC neurons (both inspiratory and expiratory) modulate cardiac parasympathetic neuronal activity [48], whilst excitatory preBötC neurons modulate sympathetic vasomotor neuronal activity [49]. Loss of neurons in the preBötC has profound effects on cardiovascular output, including respiratory sinus arrhythmia (RSA) [50] and exacerbation of hypertension [49]. Loss of both excitatory and inhibitory neurons therefore would lead to impaired respiratory entrainment in a normotensive rat, with exaggerated effects on blood pressure and heart rate in a hypertensive rat. We predict that long-term effects of the lack of entrainment would exacerbate microvascular effects of pre-existing hypertension.

Central sleep apnoea (CSA) is defined as a lack of ventilatory drive and respiratory effort during sleep, which is in stark contrast to obstructive sleep apnoea, where respiratory effort is observed during a dyspnoeic episode [51]. We observed dyspnoeic episodes that were not terminated by arousal during REM sleep in this model, the lack of arousal was replaced by a period of hypernoea to re-ventilate, which is indicative of CSA [52]. The pathophysiology of CSA may contribute to the instability of the respiratory column control of breathing during sleep by causing chronic inflammation, as intermittent hypoxia can inhibit phrenic long term facilitation [53, 54].

Characteristics from both OSA and CSA are present in this model, and the overall damage caused by their downstream effects are the same. SHRs appear to show a mixed model of sleep apnoea, containing apnoeas that are both obstructive and central in origin. With predicted damage to the recurrent laryngeal nerve by prolonged hypertension causing obstructive sleep apnoea, and loss of preBötC neurons caused by microvascular rarefaction inducing central sleep apnoea.

There is a clear link between sleep apnoea and worsening atherosclerosis in the brain, where an increase in the production of NOS [55] disrupts the vascular tone of arteries in the brain [56], leading to thickening of the artery wall specifically with an decrease of endothelial NOS (eNOS) [57], resulting in a loss of brain volume. Interestingly, where there is a decrease in upregulation of eNOS below its optimum limit, there is less regulation of blood pressure, less angiogenesis and inappropriate vascular remodelling [58], as well as the direct recruitment of activated microglia and upregulation of pro-inflammatory cytokines in the brain [59].

We saw an increase in activated microglia in both the CA1 and DG regions of the brain, areas which are associated with memory consolidation [60-62], as well as impaired long and short-term memory in the SHRs. Prolonged microglial activation is also responsible for increased damage to the blood brain barrier by causing increased permeability, which in turn causes leukocyte recruitment to the brain [63]. Some classes of leukocyte depend on VCAM1 for their migration across endothelia into the brain [64]. Once within the tissue, the leukocytes are switched by natural T killer cells, regulating their performance as reparative leukocytes. However in some cases, for example during sterile inflammation induced by sleep apnoea [29, 65], this switch does not occur and the leukocyte performs inappropriate constant damage and repair of tissue and is unable to transmigrate back into the bloodstream [66], resulting in chronic inflammation in the affected area as the cycle perpetuates. Inflammation in the brain results in white matter injury, primarily as a result of widespread demyelination [67], which may be accompanied by significant grey matter loss [68]. Myelin is particularly susceptible to death as a result of inflammation, and without the protective myelin sheath the white matter also becomes more at risk of death and therefore loss of volume [69].

We conclude that the cognitive decline present in SHRs was exacerbated due to increased neuroinflammation, that may increase microvascular rarefaction induced by prolonged hypertension [70] and chronic intermittent hypoxia [71].

## Data Availability Statement

The authors confirm the data supporting the findings of this study are available in the article and are available from the corresponding author upon request.

## Ethics Statement

All experiments involving animals were approved by the University of Warwick Animal Welfare and Ethical Review Board in accordance with the United Kingdom Animals (Scientific Procedures) Act (1986) and the EU Directive 2010/63/EU and performed under a project licence issued by the UK Home Office.

## Acknowledgments

RR performed the experiments and analysed data from all aspects of the project. RH designed and performed experiments, analysed data, and oversaw the project. Both authors contributed to the article and approved the submitted version. We would like to thank Prof. Johannes Boltze for valuable input and conversations regarding the SHR phenotype.

## Funding

This work was supported by the University of Warwick, Medical and Life Sciences Research Fund bursary, and RH is supported by a Biotechnology and Biological Sciences Research Council grant (BB/X008290/1).

## Notes

### Competing Interest Statement

The authors have declared no competing interest.

## References

1. Lim, L.L., K.W. Tham, and S.M. Fook-Chong, Obstructive sleep apnoea in Singapore: polysomnography data from a tertiary sleep disorders unit. Ann Acad Med Singap, 2008. 37(8): p. 629–36.

2. Heinzer, R., et al., Prevalence of sleep-disordered breathing in the general population: the HypnoLaus study. The Lancet. Respiratory medicine, 2015. 3(4): p. 310–318.

3. Franklin, K.A. and E. Lindberg, Obstructive sleep apnea is a common disorder in the populationa review on the epidemiology of sleep apnea. J Thorac Dis, 2015. 7(8): p. 1311–22.

4. Franklin, K.A., et al., Sleep apnoea is a common occurrence in females. European Respiratory Journal, 2013. 41(3): p. 610.

5. Zhou, B., et al., Worldwide trends in hypertension prevalence and progress in treatment and control from 1990 to 2019: a pooled analysis of 1201 population-representative studies with 104 million participants. The Lancet, 2021. 398(10304): p. 957–980.

6. Peppard, P.E., et al., Prospective Study of the Association between Sleep-Disordered Breathing and Hypertension. New England Journal of Medicine, 2000. 342(19): p. 1378–1384.

7. Young, T., et al., Population-Based Study of Sleep-Disordered Breathing as a Risk Factor for Hypertension. Archives of Internal Medicine, 1997. 157(15): p. 1746–1752.

8. Haas, D.C., et al., Age-Dependent Associations Between Sleep-Disordered Breathing and Hypertension. Circulation, 2005. 111(5): p. 614–621.

9. Wang, M., et al., Associations of genetic susceptibility and healthy lifestyle with incidence of coronary heart disease and stroke in individuals with hypertension. European Journal of Preventive Cardiology, 2022. 29(16): p. 2101–2110.

10. Sierra, C., Hypertension and the Risk of Dementia. Front Cardiovasc Med, 2020. 7: p. 5.

11. Burns, A. and S. Iliffe, Dementia. BMJ, 2009. 338: p. b75.

12. DurÁN, J., et al., Obstructive Sleep Apnea–Hypopnea and Related Clinical Features in a Population-based Sample of Subjects Aged 30 to 70 Yr. American Journal of Respiratory and Critical Care Medicine, 2001. 163(3): p. 685–689.

13. Bixler, E.O., et al., Association of hypertension and sleep-disordered breathing. Arch Intern Med, 2000. 160(15): p. 2289–95.

14. Seravalle, G. and G. Grassi, Sleep Apnea and Hypertension. High Blood Pressure & Cardiovascular Prevention, 2022. 29(1): p. 23–31.

15. Appleton, Jason P., et al., Hypercholesterolaemia and vascular dementia. Clinical Science, 2017. 131(14): p. 1561–1578.

16. Peracaula, M., et al., Endothelial Dysfunction and Cardiovascular Risk in Obstructive Sleep Apnea: A Review Article. Life (Basel), 2022. 12(4).

17. Kalaria, R.N. and T. Erkinjuntti, Small vessel disease and subcortical vascular dementia. J Clin Neurol, 2006. 2(1): p. 1–11.

18. Culebras, A. and S. Anwar, Sleep Apnea Is a Risk Factor for Stroke and Vascular Dementia. Current Neurology and Neuroscience Reports, 2018. 18(8): p. 53.

19. Rundek, T., et al., Vascular Cognitive Impairment (VCI). Neurotherapeutics, 2022. 19(1): p. 68–88.

20. Whitmer, R.A., et al., Midlife cardiovascular risk factors and risk of dementia in late life. Neurology, 2005. 64(2): p. 277–281.

21. Kaiser, D., et al., Spontaneous white matter damage, cognitive decline and neuroinflammation in middle-aged hypertensive rats: an animal model of early-stage cerebral small vessel disease. Acta Neuropathol Commun, 2014. 2: p. 169.

22. Suvila, K., et al., Age of Hypertension Onset: Overview of Research and How to Apply in Practice. Curr Hypertens Rep, 2020. 22(9): p. 68.

23. Carley, D.W., et al., Sleep-disordered respiration in phenotypically normotensive, genetically hypertensive rats. Am J Respir Crit Care Med, 2000. 162(4 Pt 1): p. 1474–9.

24. Mayor, A.H., et al., Effect of blood pressure changes on air flow dynamics in the upper airway of the decerebrate cat. Anesthesiology, 1996. 84(1): p. 128–34.

25. Garpestad, E., et al., Phenylephrine-induced hypertension acutely decreases genioglossus EMG activity in awake humans. J Appl Physiol (1985), 1992. 72(1): p. 110–5.

26. Salamone, J.A., et al., Cranial and phrenic nerve responses to changes in systemic blood pressure. J Appl Physiol Respir Environ Exerc Physiol, 1983. 55(1 Pt 1): p. 61–8.

27. Wall, M.J., et al., The Temporal Dynamics of Arc Expression Regulate Cognitive Flexibility. Neuron, 2018. 98(6): p. 1124–1132.e7.

28. McKay, L.C. and J.L. Feldman, Unilateral ablation of pre-Botzinger complex disrupts breathing during sleep but not wakefulness. Am J Respir Crit Care Med, 2008. 178(1): p. 89–95.

29. Roberts, R., et al., An Improved Model of Moderate Sleep Apnoea for Investigating Its Effect as a Comorbidity on Neurodegenerative Disease. Frontiers in Aging Neuroscience, 2022. 14.

30. Reule, S. and P.E. Drawz, Heart rate and blood pressure: any possible implications for management of hypertension? Curr Hypertens Rep, 2012. 14(6): p. 478–84.

31. Lai, C.-T., et al., Sympathetic Hyperactivity, Sleep Fragmentation, and Wake-Related Blood Pressure Surge During Late-Light Sleep in Spontaneously Hypertensive Rats. American Journal of Hypertension, 2016. 29(5): p. 590–597.

32. Ma, J., et al., Chronic brain inflammation causes a reduction in GluN2A and GluN2B subunits of NMDA receptors and an increase in the phosphorylation of mitogen-activated protein kinases in the hippocampus. Molecular Brain, 2014. 7(1): p. 33.

33. Lusis, A.J., Atherosclerosis. Nature, 2000. 407(6801): p. 233–41.

34. Austin, T.R., et al., Association of Brain Volumes and White Matter Injury With Race, Ethnicity, and Cardiovascular Risk Factors: The Multi-Ethnic Study of Atherosclerosis. J Am Heart Assoc, 2022. 11(7): p. e023159.

35. Kerkhove, D., I. Paciolla, and G. Arpino, Chapter 3 - Classification by Mechanisms of Cardiotoxicity, in Anti-Cancer Treatments and Cardiotoxicity, P. Lancellotti, J.L. Zamorano Gómez, and M. Galderisi, Editors. 2017, Academic Press: Boston. p. 13–34.

36. Yulu, S. and M.W. Joanna, Update on cerebral small vessel disease: a dynamic whole-brain disease. Stroke and Vascular Neurology, 2016. 1(3): p. 83.

37. Dingli, K., et al., Arousability in sleep apnoea/hypopnoea syndrome patients. European Respiratory Journal, 2002. 20(3): p. 733.

38. Bangash, A., et al., Obstructive Sleep Apnea and Hypertension: A Review of the Relationship and Pathogenic Association. Cureus, 2020. 12(5): p. e8241.

39. Patel, J.M.C.G., Recurrent Laryngeal Nerve Injury. StatPearls [Internet], 2023.

40. Novakovic, D. and S. MacKay, Adult obstructive sleep apnoea and the larynx. Curr Opin Otolaryngol Head Neck Surg, 2015. 23(6): p. 464–9.

41. Van den Bossche, K., et al., Natural sleep endoscopy in obstructive sleep apnea: A systematic review. Sleep Medicine Reviews, 2021. 60: p. 101534.

42. McNicholas, W.T. and D. Pevernagie, Obstructive sleep apnea: transition from pathophysiology to an integrative disease model. J Sleep Res, 2022. 31(4): p. e13616.

43. Gál, Z., et al., Investigation of the Possible Role of Tie2 Pathway and TEK Gene in Asthma and Allergic Conjunctivitis. Frontiers in Genetics, 2020. 11.

44. Brindle, N.P., P. Saharinen, and K. Alitalo, Signaling and functions of angiopoietin-1 in vascular protection. Circ Res, 2006. 98(8): p. 1014–23.

45. Augustin, H.G., et al., Control of vascular morphogenesis and homeostasis through the angiopoietin-Tie system. Nat Rev Mol Cell Biol, 2009. 10(3): p. 165–77.

46. Kong, D.H., et al., Emerging Roles of Vascular Cell Adhesion Molecule-1 (VCAM-1) in Immunological Disorders and Cancer. Int J Mol Sci, 2018. 19(4).

47. Cook-Mills, J.M., M.E. Marchese, and H. Abdala-Valencia, Vascular cell adhesion molecule-1 expression and signaling during disease: regulation by reactive oxygen species and antioxidants. Antioxid Redox Signal, 2011. 15(6): p. 1607–38.

48. Cui, Y., et al., Defining preBötzinger Complex Rhythm- and Pattern-Generating Neural Microcircuits In Vivo. Neuron, 2016. 91(3): p. 602–14.

49. Menuet, C., et al., PreBötzinger complex neurons drive respiratory modulation of blood pressure and heart rate. Elife, 2020. 9.

50. Furuya, W.I., et al., The role of glycinergic inhibition in respiratory pattern formation and cardio-respiratory coupling in rats. Curr Res Physiol, 2021. 4: p. 80–93.

51. Eckert, D.J., et al., Central sleep apnea: Pathophysiology and treatment. Chest, 2007. 131(2): p. 595–607.

52. Hanly, B.J., et al., Respiration and Abnormal Sleep in Patients with Congestive Heart Failure. CHEST, 1989. 96(3): p. 480–488.

53. Hocker, A.D., et al., The impact of inflammation on respiratory plasticity. Exp Neurol, 2017. 287(Pt 2): p. 243–253.

54. Devinney, M.J., et al., Hypoxia-induced phrenic long-term facilitation: emergent properties. Ann N Y Acad Sci, 2013. 1279: p. 143–53.

55. Valko, M., et al., Free radicals and antioxidants in normal physiological functions and human disease. The International Journal of Biochemistry & Cell Biology, 2007. 39(1): p. 44–84.

56. Matthys, K.E. and H. Bult, Nitric oxide function in atherosclerosis. Mediators Inflamm, 1997. 6(1): p. 3–21.

57. Thijssen, D.H., N.T. Cable, and D.J. Green, Impact of exercise training on arterial wall thickness in humans. Clin Sci (Lond), 2012. 122(7): p. 311–22.

58. An, L., et al., Deficiency of Endothelial Nitric Oxide Synthase (eNOS) Exacerbates Brain Damage and Cognitive Deficit in A Mouse Model of Vascular Dementia. Aging Dis, 2021. 12(3): p. 732–746.

59. Katusic, Z.S. and S.A. Austin, Endothelial nitric oxide: protector of a healthy mind. Eur Heart J, 2014. 35(14): p. 888–94.

60. Hainmueller, T. and M. Bartos, Dentate gyrus circuits for encoding, retrieval and discrimination of episodic memories. Nat Rev Neurosci, 2020. 21(3): p. 153–168.

61. Bahar, A.S., P.R. Shirvalkar, and M.L. Shapiro, Memory-Guided Learning: CA1 and CA3 Neuronal Ensembles Differentially Encode the Commonalities and Differences between Situations. The Journal of Neuroscience, 2011. 31(34): p. 12270.

62. Minhas, P.S., et al., Restoring metabolism of myeloid cells reverses cognitive decline in ageing. Nature, 2021. 590(7844): p. 122–128.

63. Zhou, H., et al., A requirement for microglial TLR4 in leukocyte recruitment into brain in response to lipopolysaccharide. J Immunol, 2006. 177(11): p. 8103–10.

64. Leick, M., et al., Leukocyte recruitment in inflammation: basic concepts and new mechanistic insights based on new models and microscopic imaging technologies. Cell Tissue Res, 2014. 355(3): p. 647–56.

65. Huber-Lang, M., J.D. Lambris, and P.A. Ward, Innate immune responses to trauma. Nature Immunology, 2018. 19(4): p. 327–341.

66. Zindel, J. and P. Kubes, DAMPs, PAMPs, and LAMPs in Immunity and Sterile Inflammation. Annual Review of Pathology: Mechanisms of Disease, 2020. 15(1): p. 493–518.

67. Rossi, S., et al., Interleukin-1β causes excitotoxic neurodegeneration and multiple sclerosis disease progression by activating the apoptotic protein p53. Molecular Neurodegeneration, 2014. 9(1): p. 56.

68. Trapp, B.D., et al., Cortical neuronal densities and cerebral white matter demyelination in multiple sclerosis: a retrospective study. Lancet Neurol, 2018. 17(10): p. 870–884.

69. Souza-Rodrigues, R.D., et al., Inflammatory response and white matter damage after microinjections of endothelin-1 into the rat striatum. Brain Research, 2008. 1200: p. 78–88.

70. Antonios, T.F., Microvascular Rarefaction in Hypertension—Reversal or Over-Correction by Treatment? American Journal of Hypertension, 2006. 19(5): p. 484–485.

71. Nanduri, J., et al., Hypoxia-inducible factors and hypertension: lessons from sleep apnea syndrome. J Mol Med (Berl), 2015. 93(5): p. 473–80.

